# Significantly Improved Mouse and Rat Genome Annotation Using Sequence Read Archive RNA-seq Data

**DOI:** 10.64898/2026.03.06.709975

**Authors:** Fan Meng, David Turner, Megan H. Hagenauer, Stanley Watson, Huda Akil

**Author notes:** Correspondence should be addressed to F.M.

## Abstract

To detect currently unannotated genes with low expression levels with high sensitivity and accuracy, we developed a new exon->gene->transcript annotation pipeline that can identify previously undetected multi-exon transcripts using large volumes of RNA-Seq data. Our pipeline incorporates three new algorithms: 1) model-based spliced exon detection, 2) exon-to-gene assignment across multiple tissue/datasets through exon community discovery, and 3) ranking top transcripts by a stepwise minimum flow procedure. The design of our pipeline allowed us to leverage hundreds of Tbases of public RNA-seq data as input to improve mouse and rat genome annotation. Using this data, our pipeline identified close to 15K and 21K unannotated genes in GENCODE M37 and ENSEMBL 114 for mouse and rat, respectively. Each species also gained over 200K predicted transcripts containing at least one new exon, although most were transcripts from GENCODE/ENSEMBL annotated genes with newly assigned exons. To make our genome annotation available for common use, we have packaged this new annotation in standard file formats for the analysis of bulk and single cell RNA-seq data (GTF, 10X genome files). We have also provided two use examples which demonstrate the utility of our newly annotated genes in functional analyses, showing that their expression can be differentially regulated in relationship to cell type and selective breeding. Due to the efficiency provided by our pipeline, we expect that as new RNA-seq data become available in the coming years it will significantly benefit rat gene/transcript annotation, eventually enabling us to approach the target of complete gene and transcript annotation.

## Introduction

Mice (*Mus musculus*) and rats (*Rattus norvegicus*) are widely used model organisms for understanding human biology and diseases. While their gene and transcriptome annotation has significantly improved with the release of GENCODE M36 for mouse and ENSEMBL 114 for rat, the large difference in mouse and rat gene counts (78K vs 44K) suggests that at least the rat gene annotation is far from completion.

Better annotation of mouse and rat genomes will improve gene-level analyses in rodent disease models and will also help identify genes and transcripts that differ in rodents from humans and may be human or primate-specific. Understanding what makes us human is a perennial topic in biology but past studies at the gene level were often hampered by incomplete gene annotations (Gradnigo et al. 2016; Weisman et al. 2020; Weisman et al. 2022).

Most of the currently unannotated genes are likely to be long non-coding RNA (lncRNA) genes. Compared to protein-coding genes, lncRNA genes are typically expressed at a much lower level in tissues. They also exhibit higher tissue and cell type specificity. Unlike protein coding sequences, lncRNA sequences evolve rapidly across mammalian species although their secondary structure and relative genomic positions, *i*.*e*., synteny, are conserved better than their sequences (Mattick et al. 2023).

lncRNAs are known to be involved in many different biological processes (Rinn and Chang 2020; Statello et al. 2021; Mattick et al. 2023), such as epigenetic regulation through dosage compensation (Lee 2012) and genomic imprinting (Bartolomei et al. 1991), transcription control through chromatin modification (Rinn et al. 2007) and transcription factor binding (Kino et al. 2010), nuclear body assembly (Clemson et al. 2009), splicing regulation (Tripathi et al. 2010), as well as post-transcriptional regulation by acting as miRNA “sponges” (Cesana et al. 2011) or mRNA stability control (Lee et al. 2016).

Given the importance of more complete gene/transcript annotation and the potential functional significance of lncRNA, we decided to utilize mouse and rat Illumina short-read RNA-seq data in the public database Sequence Read Archive (SRA) to improve gene/transcript annotation for mice and rats. Several algorithms have already been developed for this purpose. For example, the widely-used *StringTie* (Pertea et al. 2015; Kovaka et al. 2019) or the earlier *Cufflinks* (Trapnell et al. 2012) algorithms were developed in the early years of RNA-seq to help researchers to identify unannotated transcripts or “dark matter” for their targeted tissues and diseases, as many genes were not annotated at that time. These two algorithms also enabled large scale RNA-seq based transcript annotation projects such as *miTranscriptome* (Iyer et al. 2015) and *CHESS* (Pertea et al. 2018) that provided more extensive gene and transcript annotation for the human genome than GENCODE/ENSEMBL at the same periods.

We found that existing algorithms were not ideal for identifying currently unannotated genes, which are likely to be expressed at lower levels than known genes. For these genes, transcript annotations based on individual samples would be expected to have low sensitivity for exons that are only expressed in a small fraction of cells. For example, when we started our project, we used *Stringtie2*, which is the most widely used algorithm for RNA-seq data based new transcript annotation. However, we soon realized that individual RNA-seq samples often lacked sufficient coverage for multi-exon transcripts with low expression levels, causing reads to appear to form “single exon” transcription fragments rather than connected multi-exon transcripts.

Merging RNA-seq data from several hundred samples from the same tissue and developmental stage can potentially address this issue, but existing gene and transcript annotation algorithms based on RNA-seq data are not designed to process tera base (Tbase) level data. During our initial attempts using *Stringtie2*, we found that when we merged RNA-seq data from dozens or hundreds of samples for the purpose of detecting low expression multi-exon transcripts, intron reads and random transcription noise accumulated and led to continuous transcripts spanning multiple exons and introns, or even spanning different genes for highly transcribed regions. Limitations to *StringTie* were also noted by researchers in the GENCODE Capture Long Read Sequence (CLS) annotation project, where the algorithm generated overly long 5’ extensions (Lagarde et al. 2017), likely due to overlapping transcription noise.

As a result, we decided to develop a new exon->gene->transcript annotation pipeline using a totally different approach (**Fig. 1**). Our method focuses on spliced exons for higher accuracy and sensitivity, *as real splicing signals should accumulate while noise should flatten out when RNA-seq data from many samples are merged*. Our pipeline incorporates three new algorithms: 1) model-based spliced exon detection, 2) exon-to-gene assignment across multiple tissue/datasets through exon community discovery, and 3) ranking top transcripts by a stepwise minimum flow procedure. Importantly, our pipeline can handle the hundreds of Tbases of RNA-seq data from mice and rats that are now available in SRA.

**Fig 1.**
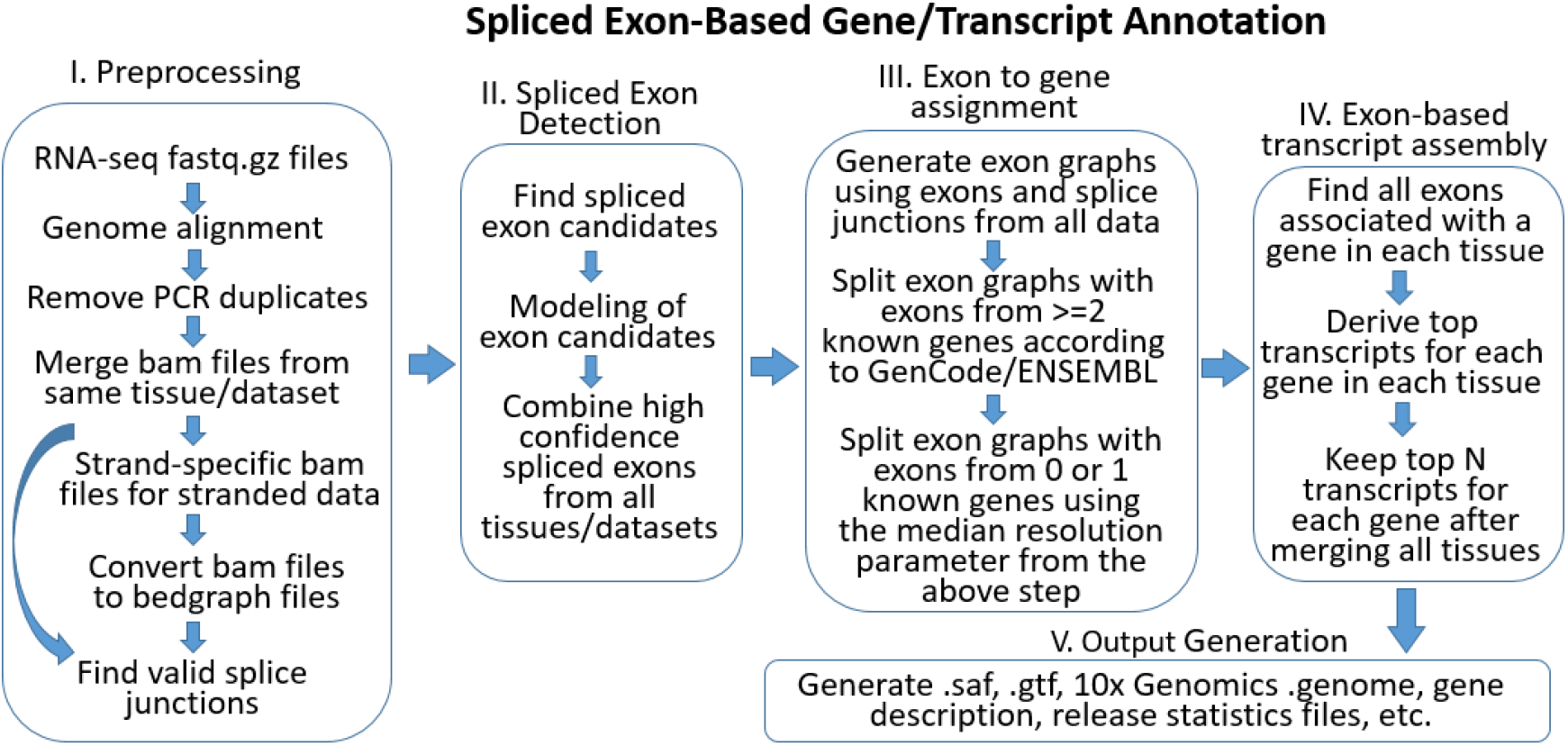
A new pipeline for multi-exon gene/transcript annotation using large-scale RNA-seq data (>Tb) and spliced exon detection. To identify currently unannotated genes, we designed a pipeline that was optimized for the detection of low-level expressed transcripts using merged RNA-seq data. For optimal coverage, we used several hundred samples from the same tissue and development stage, requiring the generation of an algorithm that could process tera base (Tb) level RNA-seq data. Our method focuses on spliced exons as real splicing signals should accumulate while noise should flatten out when RNA-seq data from many samples are merged. Following RNA-seq data preprocessing (step 1), our pipeline incorporates three new algorithms: model-based spliced exon detection (step 2), exon-to-gene assignment across multiple tissue/datasets through exon community discovery (step 3), and top transcripts ranking by a stepwise minimum flow procedure (step 4). The algorithm then generates output (step 5) in file formats commonly used for genomic data analysis.

We applied our pipeline for the detection of unannotated exons and genes for mice and rats. The new exon, transcript and gene annotations identified by our pipeline were merged with annotations in GENCODE M37 and ENSEMBL 114 for mouse and rat, respectively to generate standard genome reference files necessary for bulk RNA-seq analysis and single cell/nucleus RNA-seq (scRNA-Seq, snRNA-Seq) analysis. Finally, we used these files to demonstrate the insight provided by our new annotation in two use case examples (bulk RNA-seq and snRNA-Seq).

## Methods

All computing tasks were completed on a local computer cluster with 418 hyperthreading capable cores, 9.6TB physical memory and 1.5 PB high-speed storage space. It took about 6 months to finish the mouse and rat annotation from scratch using our pipeline optimized for human data based on GENCODE 39.

### RNA-seq data used for novel spliced exon detection

To detect low-level expressed transcripts, we used merged RNA-seq data representing hundreds of samples from the same tissues and development stages for both mice and rats. As SRA hosts many more RNA-seq datasets derived from mice than rats, we used paired mouse RNA-seq datasets with at least 10 samples and having >50 Gbases of data, but included all rat RNA-Seq datasets having >10 Gbases. We excluded some mouse datasets from over-represented tissues such as liver, lung, blood, whole brain, bone marrow, etc., to keep physical memory usage < 2TB at the *Porticullis-*based splice site filtering stage (Mapleson et al. 2018), which is the most memory-demanding stage of the pipeline. Studies involving cancer/tumors, unrelated species, or non-RNA-seq data that passed the initial keyword-based filtering were also removed before data download. The mouse and rat data were downloaded in March and April of 2025. A total of 821 mouse datasets (∼ 400 TBases) and 1673 rat datasets (∼ 200 TBases) were grouped into 184 and 223 tissue-development stage groups for spliced exon discovery and transcript assembly.

### Step 1: RNA-seq data preprocessing

The RNA-seq data was preprocessed using an analysis pipeline built from existing programs such as *fastp* (Chen et al. 2018; Chen 2023), the *STAR aligner* (Dobin et al. 2013), *SAMtools* (Li et al. 2009), *BEDTools* (Quinlan and Hall 2010) and *Portcullis* (Mapleson et al. 2018). Genome alignment was performed using standard ENCODE parameters, but we only allowed a 2% error rate to reduce noise. Identically mapped paired-end or single-end reads (PCR duplicates) in each RNA-seq sample were removed to reduce the effect of uneven PCR amplification before merging bam files from the same tissue-development stage. High confidence splice junctions in the merged bam files were identified using the *Portcullis* program (Mapleson et al. 2018). Typically, 30%-40% of the STAR aligner detected splice junctions passed the Portcullis filters. If a merged bam file contained paired strand-specific reads, they were further processed to generate strand-specific bam files to take advantage of their significantly better signal-to-noise ratio. All bam files, including the original bam files that contain alignments to both strands of the genome, were converted to bedgraph format for spliced exon detection at the *Portcullis* filtered splice sites.

### Step 2: Spliced exon detection

Our spliced exon detection method is based on distinctive patterns of genome-aligned read coverage shown by known spliced exons when aligned RNA-seq reads are displayed in a genome browser such as *Integrative Genomics Viewer (IGV)* (Robinson et al. 2011; Thorvaldsdottir et al. 2013). Consequently, we can turn exon detection into a model fitting problem (Fig. 2).

**Fig 2.**
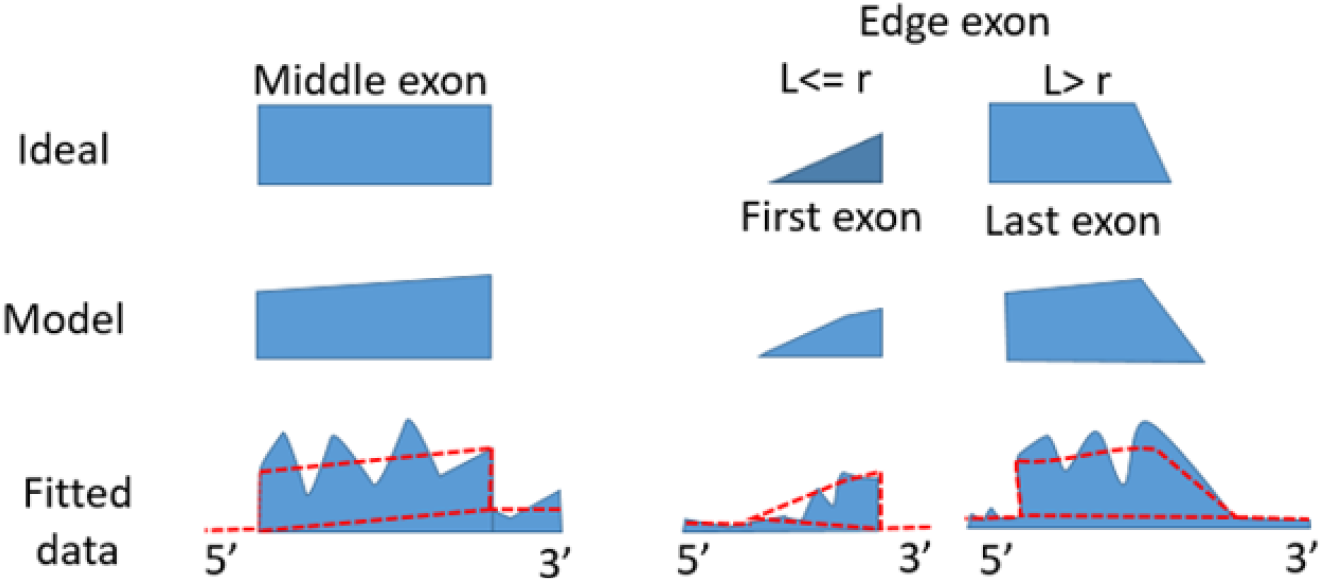
Exon signal models for spliced exons.

The “Ideal” row in Fig. 2 illustrates the expected RNA-seq read coverage signal patterns for different type of exons without real world complications. We call the first or the last exon in a transcript “edge exon” and the remaining exons “middle exon”. Note that when exon length *L* is greater than the read length *r*, the base length of the slope region of edge exons should be *r-1* due to the lowering probability of reads hitting the regions closer to the exon boundary. Since most data were from polyA-selected libraries, the actual models are listed in the “Model” row to account for 3’ bias. We used trapezoids with two vertical edges to model middle exon signals. Quadrilaterals with one vertical edge were used to model edge exons regardless of the *L* vs. *r* relationship. This is because the length of the edge exon is not known beforehand and, even if exon length is shorter than read length, using a quadrilateral with a right angle rather than a right triangle does not influence the results very much due to sequencing noises. This is also one of the reasons that we did not enforce the *r-1* slope region length for edge exons, besides potential multiple transcription start/termination sites.

The key issue for edge exons is to determine the boundary for the non-spliced end. We started from 5 boundary locations, estimated at 5 background signal cutoffs, within 3 kb from the splice site and search for the closest (to the splice site) signal valley that <= ½ of signal at the splice site using the R *CKmeans*.*1d*.*dp* package. The goal was to obtain a boundary estimation on the conservative side (shorter) but not remove too much read signal. This also meant that we could not determine the exact start/termination sites of transcripts, although we could identify the first and the last exons for most transcripts.

We used the R *nlstools* package to fit the multi-segment line models for middle and edge exons, including background regions before and after the exon. The background noise level for a candidate exon is determined by the bottom line between two background points on both sides of the candidate exon (“Fitted data” row in **Fig. 2**). Models with fitting residuals <60%, i.e., read signal >=40%, in the expected exon signal areas were considered valid exons. This is because *requiring the model area to be filled with a minimum amount of read signal removes spurious splicing signals from background noise or the presence of introns*. To ensure the detection of major exon splicing variants, we required signal-to-background ratio at the spliced edge >=1.2 as well as >= 20 junction-spanning reads in each tissue-development group. These parameters were optimized using the human GTEx V8 dataset together with GENCODE 39 to achieve ∼ 85% known exon detection.

After exon discovery in each tissue-development group, we merged results from all tissues and selected all middle exons that passed the above-mentioned exon-intron interface and model fitting thresholds in any single group as detected middle exons. Since we could not accurately determine the exact location of the non-spliced end of edge exons, we selected one edge exon from multiple edge exons sharing the same splice site and used the start-end location of the selected edge exon to replace all edge exons having the same splice site in different tissue-development groups. The selection was based on the edge exon length probability distribution derived from known mouse or rat edge exons in GENCODE M37 and ENSEMBL 114, respectively. An edge exon with the highest length probability and that was shorter than other edge exons with identical length probability, when it existed, was selected.

As a result, our annotation only kept one edge exon for every unique splice site. This is different from GENCODE/EMSEMBL annotations, in which there are often multiple edge exons sharing the same splice site but with different non-spliced ends. Our approach may miss multiple transcription start/termination sites for the same edge exon but is an acceptable compromise given the noisy nature of the non-spliced site location signal in RNA-seq data. A technical advantage of only keeping one edge exon for each splice site is that it often significantly reduces the computation time needed at the transcript assembly step.

### Step 3: Exon to gene assignment

To assign new exons to genes using data from all of our chosen RNA-seq datasets, we first created connected exon graphs for all known and new exons using *Portcullis* filtered exon junctions. An unexpected issue with large volumes of RNA-seq data from different tissue-development stage datasets was that a connected exon graph often contained more than one known gene. While a small fraction of them probably reflected GENCODE/ENSEMBL annotation issues, most of them were likely caused by spurious splicing or read-through events. Since exons from the same gene should have a higher number of junction spanning reads between each other than exons from other genes, *we assert that assigning exons to genes in an exon graph is equivalent to the community discovery problem for directed graphs*.

We found that the Leiden algorithm (Traag et al. 2019) provided the best results and computational efficiency when tested on exon graphs containing exons from 2 or more known genes in GENCODE 39 with the GTEx V8 dataset. For each exon graph containing exons from two or more known genes, and frequently also new exons, we gradually increased the Leiden clustering resolution parameter (LCRP) from 0.0001 until all exons from different known genes were properly separated. Sometimes all new exons, when they existed, were assigned to known genes at the same time. If some new exons were separated from known genes into new cluster(s), the new cluster(s) with >=2 exons were defined as new gene(s). After all exon graphs containing exons from 2 or more known genes were processed, we use the median LCRP value derived from all such cases to process exon graphs containing exons from 1 or 0 known genes. That approach allowed us to take advantage of curated gene definitions in GENCODE/ENSEMBL in the process of assigning new exons to known genes as well as to new genes.

### Step 4: Exon-based transcript assembly

After the above exon-to-gene assignment step, we had a clean exon graph describing the relationships among all exons from different tissue-development stage groups for the same gene. Our exon graph-based transcript assembly was then performed for each tissue-development stage group based on group-specific exons and their relationships for finding top transcripts in each group. To speed-up the top N transcript identification, we first used Leiden clustering to decompose complex exon graphs for genes with many first/last exons into the largest possible number of subclusters, under the condition that each subcluster had at least 1 first exon and 1 last exon, by gradually increasing LCRP. This was because *top transcripts should have higher junction-spanning reads along their 5’ to 3’ paths* and thus can be separated from low abundance transcripts through directed graph clustering.

The R *igraph* package *all_simple_paths* function was then used to identify all possible transcripts between any pair of first and last exons for selecting transcripts with highest abundance levels. The abundance for each predicted transcript from the same gene was ranked using a stepwise minimum flow approach: all predicted transcripts were first ranked based on the minimum flow, *i*.*e*., the splice junction with lowest number of read counts. If several transcripts had the same minimum flow, they were further ranked using the second lowest flow. If a shorter transcript ran out of junctions, it was ranked above all longer transcripts in the same group. This process was repeated until no transcript had the same order in the top N transcripts that we wanted to keep. Compared to all previous transcript ranking approaches for RNA-seq data based transcript assembly (Pertea et al. 2015), our approach is closer to the nature of mRNA precursor processing: a partially processed precursor with higher concentration will have a better chance to finish the next splicing step.

### Step 5: Output generation

Our transcript assembly output only kept transcripts with at least one new exon not included in current GENCODE/ENSEMBL and that passed length (transcript length >=500 bp) and gene expression abundance cutoffs (average splice junction read depth for a gene from all samples >=80). We only kept unannotated genes that produced at least one >=500 bp transcripts according to the recent lncRNA definition (Mattick et al. 2023). The average total splice junction read count for all splice junctions in a gene, after removing PCR duplicates within each sample, was used to ensure the annotated genes were expressed in multiple subjects and tissues to avoid the inclusion of transcription noise. We removed all predicted transcripts that did not alter known gene structure, including fragments of known transcripts, only with exon skipping, or identical to known transcripts.

These currently unannotated genes and transcripts, the majority being transcripts containing unannotated exons from known genes, were then merged with gene and transcript annotations from GENCODE M37 for mouse and ENSEMBL 114 for rat for downstream analysis. We generated standard GTF files for bulk RNA-seq analysis as well as 10x genome files for single cell/nucleus RNA-seq analysis.

#### Comparison of results from our pipeline with GENCODE M37

To make results from our pipeline comparable with GENCODE M37, we needed to make two adjustments: 1) since our pipeline only annotated genes containing splice sites, we only compared our results to multiexon GENCODE M37 genes, 2) we used a less stringent >=200 bp filter for transcripts from our pipeline to match the lack of length filter used by GENCODE M37. We considered a GENCODE exon to be detected if there was one exact splice site overlap with our predicted exon(s) on the same strand. Similarly, a gene was considered as detected as long as at least one of its exons is detected.

#### Use case examples: Single cell and bulk RNA-seq data analysis

To demonstrate utility for single cell RNA-Seq analysis, we re-analyzed raw data from *GSE243413* (Li et al. 2024) using 10x genome files derived from our mouse gene annotation to examine the role of new genes we annotated as cell type marker genes in the different retina cell types annotated by the original authors. To preprocess the data, we used *10x Genomics CellRanger* 9.0.1, followed by the *Seurat V5* pipeline (Hao et al. 2024). We used the *FindAllMarkers* function in the *Seurat* package for cell type marker discovery among different retina cell types. To reduce the influence of cell number differences among different cell types, we down-sampled cell counts to 300 for large cell types. To decrease the impact of random sampling, we performed 100 runs of *FindAllMarkers*, each with a different random sampling seed. The final results were averaged across runs and we also counted the number of times a gene passed the 2-fold enrichment threshold at *padj*<0.05 in 100 runs. Only genes that showed up at least 50 times were kept as potential cell type markers.

To demonstrate utility for bulk RNA-Seq analysis, we re-analyzed the raw hippocampal RNA-seq data described in a recent paper ((Hebda-Bauer et al. 2025): *GSE225744*) using the updated rat gene annotation that we generated. This dataset characterizes the brains of rats that have been selectively bred to show low locomotor response to a novel environment (bred Low Responders or bLRs) or high locomotor response to a novel environment (bred High Responders or bHRs), which have replicated, robust differences in hippocampal gene expression. To re-analyze the data, we followed the analysis procedure provided in the code release accompanying the paper (https://github.com/hagenaue/NIDA_bLRvsbHR_F2Cross_HC_RNASeq/tree/main/F0), which uses the *limma-voom* pipeline for differential expression analysis of bulk RNA-seq data (Ritchie et al. 2015).

## Results

### Improved mouse and rat gene/transcript annotation

Our annotation increased the number of identified mouse genes by close to 15K (18.6%) beyond GENCODE M37, which already included almost 22K new genes from the GENCODE CLS project (Kaur et al. 2024). The improvement for rat gene annotation was more significant: we added close to 21K genes beyond the ENSEMBL 114 rat gene annotation, representing a 48.3% increase in identified rat genes due to inadequate annotation for the rat genome in the last decade. **Table 1** provides the summary statistics for genes, exons and transcripts identified with our annotation.

**Table 1.** Summary of updated mouse and rat gene annotation*.

|  | Gene<br>Total | Exon<br>Total | New<br>Exon in<br>Known<br>Gene | New Exon<br>in New<br>Gene | Tx Total | New Tx<br>in<br>Known<br>Gene | New Tx in<br>New Gene |
| --- | --- | --- | --- | --- | --- | --- | --- |
| Updated mouse | 92888 | 876213 | 119053<br>(30499) | 64098<br>(14571) | 517126 | 195111<br>(30499) | 43646<br>(14571) |
| GENCODE M37 | 78317 | 693062 | / | / | 278369 | / | / |
| Updated rat | 64293 | 570908 | 89166<br>(22355) | 112400<br>(20933) | 346883 | 157278<br>(22355) | 94133<br>(20933) |
| ENSEMBL114 rat | 43360 | 369342 | / | / | 95472 | / | / |

\*Numbers in parentheses are the corresponding gene counts.

An unexpected result was that most of the new transcripts that we predicted are actually transcripts from known genes but with at least one new spliced exon annotated by our pipeline, rather than transcripts from new genes as we originally expected. Around 30K GENCODE M37 genes and 20K ENSEMBL114 rat genes were assigned one or more new exons by our pipeline. These new exons can potentially alter transcription regulation, transcript stability as well as altered open reading frames (ORFs). While additional studies are needed to understand their exact functional consequences, our work suggests that many known genes may still not have full exon annotations.

### Comparison of annotation results from different methods

We compared the results from the whole genome annotation generated by our pipeline using ∼60K mouse short-read samples to the annotation generated by 1) the GENCODE CLS project using long read sequencing of captured transcripts from likely transcribed genomic regions from only a couple dozen samples and 2) the standard GENCODE pipeline without CLS contribution, which is defined as stable gene ids in M35 but not in M36.

**Table 2** is a summary of the comparison results. It shows that our pipeline detected about 90% of M37 CLS as well as non-CLS multiexon genes. It caught around 85% of exons in all M37 genes although the detection rate for CLS genes is more than 5% lower than that of non-CLS genes. The average exon count per transcript is significantly higher from our pipeline but that may be partly caused by the fact that we only keep transcripts with at least one new exon for all M37 genes.

**Table 2.** Comparison of different annotation methods for multi-exon genes*.

|  | M37 Gene | Detected Gene (%) | M37 Exon | Detected Exon (%) | M37 Tx Exon/Tx | M37 Gene New Tx Exon/Tx* |
| --- | --- | --- | --- | --- | --- | --- |
| M37 CLS multiexon | 19849 | 18004 (90.7%) | 119291 | 99036 (83.0%) | 2.96±1.22 | 4.43±2.38 |
| M37 non-CLS multiexon | 34103 | 30308 (88.9%) | 546545 | 482871 (88.3%) | 5.61±6.10 | 12.88±16.36 |

\* Based on $\geq 200$ bp predicted transcripts from our pipeline with all M37 multiexon genes and transcripts.

In total, our pipeline failed to detect 5,640 M37 multiexon genes and 83,929 of their exons from both CLS and traditional GENCODE annotation. Naturally, we also missed 24,365 single exon genes and their exons in M37, with 1,768 of them from the CLS project.

On the other hand, even at the transcript length >=500 cutoff, our pipeline found 14,571 new multiexon genes not annotated by CLS or traditional method in GENCODE M37. Among the 183,151 new exons that we annotated, 23,576 of them overlap with 36,071 exons in GENCODE M37. The vast majority of the 23,576 exons share one splice site with GENCODE M37 exon(s) but there are several hundred of them do not share any splice site with the overlapping M37 exons on the same strand. As a result, neither CLS nor traditional GENCODE annotation detected 159,575 (183,151-23,576) exons found by our pipeline.

### Use case examples: Regulated expression of newly annotated genes

Almost all mouse and rat tissue-development groups contributed new transcripts with at least 500 bp. Here we provide two examples demonstrating that the expression of some newly annotated genes can be readily detected in different tissues and many of them are under regulated expression.

#### 1. Cell marker genes in mouse retina

We re-analyzed raw data from *GSE243413* (Li et al. 2024) to examine the role of new genes we annotated as cell type marker genes in different retina cell types. **Table 3** is a summary of the distribution of GENCODE M37 genes and our newly identified genes in different retina cell types at the10-fold enrichment threshold.

**Table 3.** Mouse retina cell type marker genes with >= 10-fold enrichment.

| Retina Cell Type | GCM37 Gene | New Gene | % of New Gene |
| --- | --- | --- | --- |
| rod bipolar cell | 135 | 29 | 17.7 |
| type 7 cone bipolar cell (sensu Mus) | 51 | 10 | 16.4 |
| type 5a cone bipolar cell | 59 | 11 | 15.7 |
| type 8 cone bipolar cell (sensu Mus) | 48 | 8 | 14.3 |
| type 6 cone bipolar cell (sensu Mus) | 49 | 7 | 12.5 |
| type 9 cone bipolar cell (sensu Mus) | 77 | 10 | 11.5 |
| type 1 cone bipolar cell (sensu Mus) | 61 | 7 | 10.3 |
| type 3a cone bipolar cell | 75 | 8 | 9.6 |
| type 5 cone bipolar cell (sensu Mus) | 41 | 4 | 8.9 |
| type 3b cone bipolar cell | 124 | 12 | 8.8 |
| retina horizontal cell | 475 | 45 | 8.7 |
| type 4 cone bipolar cell (sensu Mus) | 148 | 14 | 8.6 |
| retinal cone cell | 175 | 16 | 8.4 |
| glycinergic amacrine cell | 301 | 23 | 7.1 |
| type 2 cone bipolar cell (sensu Mus) | 111 | 8 | 6.7 |
| amacrine cell | 212 | 10 | 4.5 |
| retinal rod cell | 223 | 10 | 4.3 |
| Mueller cell | 887 | 33 | 3.6 |
| GABAergic amacrine cell | 411 | 12 | 2.8 |
| endothelial cell | 1253 | 36 | 2.8 |
| microglial cell | 1055 | 16 | 1.5 |
| retinal ganglion cell | 570 | 8 | 1.4 |
| pericyte | 647 | 8 | 1.2 |
| retinal pigment epithelial cell | 413 | 3 | 0.7 |

While the overall percentage of new genes that were identified as cell type markers is not high, they exhibit highly uneven distribution across different cell types. The bipolar cell types appeared to have a higher proportion of new genes, showing a >=10-fold enrichment compared to other cell types. We had hypothesized that the new genes, due to their harder to detect nature, might play a more significant role for different retinal ganglion cell subtypes since they showed more rapid evolution and diversity across species than bipolar cells (Baden 2024). Unfortunately, this mouse retina single cell dataset only captured several hundred ganglion cells and did not provide subtype annotation among them. In this dataset there was more bipolar cell diversity than other retinal cell types and the new genes we annotated seem to play a more significant role in the separation of the different bipolar cell types. In general, the higher propensity of new genes being retinal bipolar cell markers and their highly uneven distribution across different retina cell types suggest that these genes are not randomly expressed, although their exact roles in different cell types need further investigation.

#### 2. Differential expression in the hippocampus in a selectively-bred rat model of internalizing vs. externalizing behaviors (bLR vs. bHR)

To determine whether our newly-annotated genes would show differential expression in bHR/bLR models, we re-analyzed the raw bHR/bLR hippocampal RNA-seq data described in a recent paper ((Hebda-Bauer et al. 2025): *GSE225744*) using the updated rat gene annotation that we generated. **Table 4** is a summary of differential expression analysis results across different gene biotypes, as annotated by ENSEMBL 114, at the *padj*<0.1 and *abs(log2FC)*>log2(1.5) cutoff.. The biotype category “unassigned” is used for new genes that we annotated, which should predominantly be lncRNA genes.

**Table 4.**
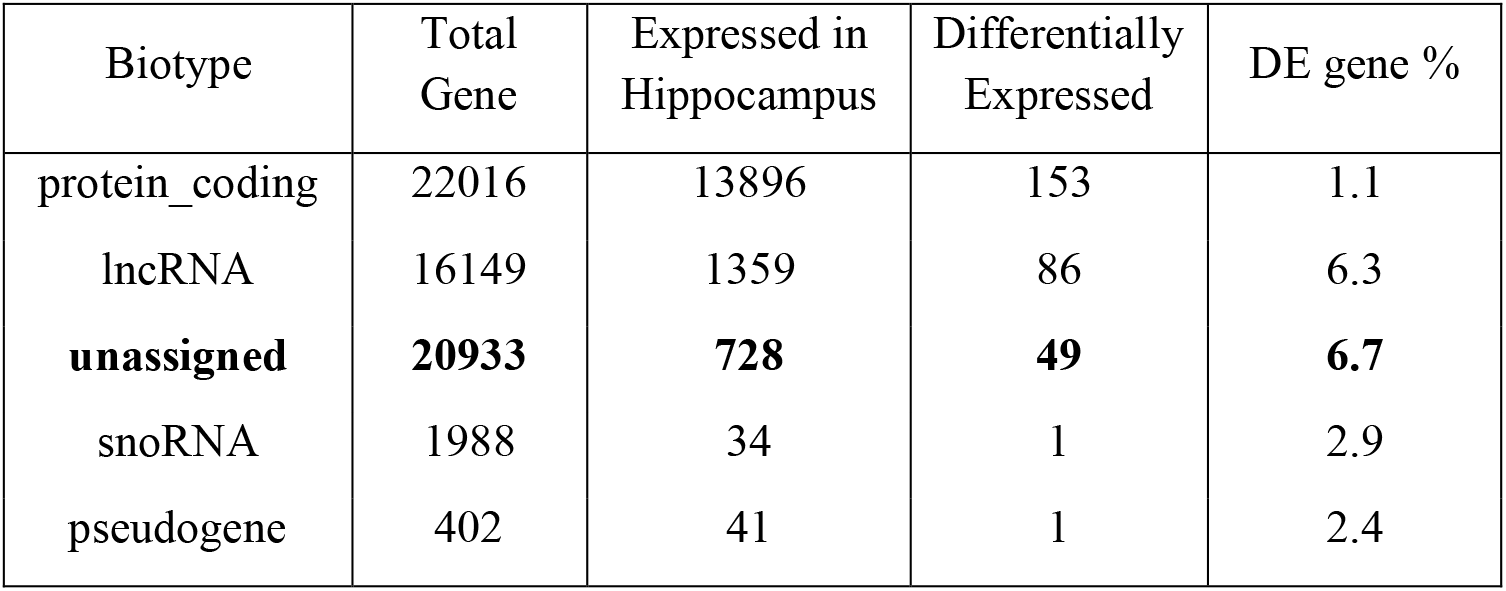
Differential expression in hippocampus of bLR vs bHR rat behavior model.

Among the 728 new rat genes that we annotated and expressed in the hippocampus, 49 are differentially expressed in the bLR vs bHR rat model comparison. The last column of **Table 4** shows the percentage of differentially expressed genes among detected hippocampal genes in each biotype. The fact that the new genes that we identified exhibited the highest percentage of differential expression suggests that they are indeed under regulated expression at a level similar to ENSEMBL 114 annotated rat lncRNA genes. The low fraction of new genes that are expressed in the hippocampus compared to known lncRNA genes probably reflects the overall lower expression level of these previously unannotated genes and their potentially higher tissue specificity.

## Discussion

Our new RNA-seq data-based gene and transcript annotation pipeline, in combination with several hundred TBases SRA RNA-seq data, significantly improved mouse and rat genome annotation. While our work increased mouse and rat gene counts to around 93K and 64K, respectively, the gene count gap between mouse and rat was only reduced slightly. The RNA-seq data size difference used for mouse and rat in this work was probably the main reason. We were able to collect bout 400 Tbases mouse data but only 200 Tbases rat data, despite using a more liberal >10 Gbases minimum dataset size cutoff for rat but > 50GBases for mouse. If public mouse and rat RNA-seq data size continues to grow at the current pace, i.e., more than doubling every two years, we can expect that the rat annotated gene count may catch up with the current mouse gene count in about two years.

Compared to new mouse gene annotations by GENCODE M37, which included 22K new genes from the GENCODE CLS project, our pipeline missed around 10% of M37 genes and around 15% of their exons for multiexon genes when we examined all transcripts with lengths >=200 bp instead of using our final >=500 bp threshold. Conceivably, the CLS procedure has much higher sensitivity in detecting all exons in a long transcript once it is captured for sequencing. In contrast, short-read RNA-seq is known to have lower sensitivity for detecting 5’ end exons as well as exons with low GC content. The second factor is that embryonic mouse tissue samples seemed to account for 25% of the total tissue samples used in the CLS project (Kaur et al. 2024). In contrast, mouse data used in our current study only included less than 2% embryonic and neonatal samples and thus was likely to miss many genes not expressed in adulthood. Another obvious advantage of the CLS project is that transcript annotations from long-read RNA-seq should be more reliable than our short-read derived annotations.

On the other hand, our approach detected close to 15K more genes and 160K more spliced exons than the full GENCODE M37 annotation. This is not surprising since the public domain short-read RNA-seq data has several orders of magnitude higher sequencing depth on most mammalian genomes than the cDNA/EST sequences that traditional GENCODE/ENSEMBL annotation mainly relies on. In addition, the mouse CLS project 1) targeted 2.8% of mouse genomic region that are likely transcribed based on multiple criteria 2) performed RNA capture and long-read sequencing only on a couple dozen or so mouse tissue samples (Kaur et al. 2024). Our approach used RNA-seq data from the whole mouse genome from 184 tissue-development stage groups derived from over 60K samples. Our results suggest that spliced genes can be transcribed from genomic regions not thought to have high transcription probability. In addition, while it is amazing that the CLS project can detect 22K new genes over M35 with just two dozen or so samples, it has limited scalability. Our pipeline enables us to use a highly scalable brute-force approach with public domain data at a much lower cost to improve multiexon mouse gene annotation.

It can be expected that while long-read RNA-seq data will increase significantly in the coming years, sample diversity and data volume from short-read RNA-seq will still offer significant advantages in the next several years. However, while the capture long read sequencing approach may not be feasible for a large number of samples across the whole genome due to cost limitations, standard long-read sequence data should be incorporated in future pipelines for better transcript assembly as well as detection of exons missed by short-read RNA-seq. Future data collection efforts should also focus more on data from developmental tissues/cells, which accounts for <2% of mouse data in this work, as well as under-represented tissues/cells.

The fact that the GENCODE CLS project could add 22K mouse genes in the GENCODE M36 release and our work added another 15K mouse genes to GENCODE M37 suggests that a central goal of the Human Genome Project, which is to determine the location of all human genes (Amaral et al. 2023), may still be a challenge after more than 20 years. At least in mouse and rat, the number of genes and exons keep increasing, although slower for mouse now, with adoption of long-read sequencing as well as larger volumes of short-read data. It seems that simply relying on existing data and solutions, such as the pipeline described in this paper or the GENCODE CLS pipeline, may not help us to achieve complete gene annotation any time soon.

A key limitation of existing short/long-read based gene and transcript annotation solutions is their “shallowness”, i.e., they mainly rely on local RNA-seq data to derive annotations for the specific input data. Other helpful information, such as nearby transcription start sites, polyA signals, translation start sites, splicing sites, neighboring gene expression level, SNP alleles, etc., are not utilized in the annotation pipeline. With the rapid development of deep learning models, it is conceivable that we could utilize all existing genome sequences, genomic elements annotation, RNA-seq data from species, tissues, cell types and developmental stages to build a foundation model to capture complex relationships underlying transcription. Such a deep learning model will enable us to have more complete annotation of genes and exons with shallow RNA-seq data.

Although lncRNA genes are involved in various biological processes, whether each of these newly annotated or known lncRNA genes play a functional role in mouse or rat is still an open question. Given the high frequency of random sequences that contain gene forming segments (Weisman 2022), it is conceivable that some new genomic regions are transcribed, spliced or even translated due to new genomic mutations, structural changes or recombination in each new individual. However, only genes that offer functional advantages will be passed across multiple generations and exhibit increased frequency in a population. As a result, only genes that can be detected in multiple subjects have the potential of being functional. Transcripts that can only be detected in one subject or even a few generations may still under regulated expression but have yet to be functionally selected and thus may not be functional. Since our pipeline requires 1) each exon splice site to have >= 20 exon-spanning reads in each tissue-development group and 2) each gene to have average accumulated splice junction reads per junction >=80, after removing PCR duplicates in individual samples as well as removing tumor/cancer samples, we suspect that the vast majority of genes annotated by our pipeline should already be functionally selected.

We excluded tumor/cancer samples in this work since we aim at the annotation of normal mouse and rat genome. The clonal expansion of tumors is known to be a dynamic process where a single mutated cell gains a fitness advantage, proliferates, and evolves into a diverse population. During this progression, gene expression changes reflect cell’s adaptation to metabolic stress, immune evasion and spatial constraints (Cho et al. 2024; Pak et al. 2026). This Darwinian selection process conceivably involves the transcription of new genomic locations due to mutation/structural change besides known genes thus likely lead to novel transcripts absent in normal tissues. If these novel transcripts are functionally important for clonal expansion, they should be easy to detect in tumor samples assuming a limited number of mechanisms that will enable key tumor phenotypes, such as immune evasion and treatment resistance. Consequently, identifying novel transcripts involved in clonal expansion of tumors may provide additional understanding of their underlying mechanisms and potential new highly specific therapeutic approaches.

In summary, our pipeline significantly improves mouse and rat gene annotation by taking advantage of the public domain short-read RNA-seq data. It is highly scalable and can be applied to other less annotated genomes and tumors with rich RNA-seq data. Our solution can be further improved by the incorporation of deep learning methods as well as long-read RNA-seq data.

## Supporting information

mouse_SRA_GENCODE_M37.gtf

rat_SRA_ENSEMBL114.gtf

## References

Amaral P, Carbonell-Sala S, De La Vega FM, Faial T, Frankish A, Gingeras T, Guigo R, Harrow JL, Hatzigeorgiou AG, Johnson R et al. 2023. The status of the human gene catalogue. Nature 622: 41–47.

Baden T. 2024. The vertebrate retina: a window into the evolution of computation in the brain. Curr Opin Behav Sci 57.

Bartolomei MS, Zemel S, Tilghman SM. 1991. Parental imprinting of the mouse H19 gene. Nature 351: 153–155.

Cesana M, Cacchiarelli D, Legnini I, Santini T, Sthandier O, Chinappi M, Tramontano A, Bozzoni I. 2011. A Long Noncoding RNA Controls Muscle Differentiation by Functioning as a Competing Endogenous RNA (vol 147, pg 358, 2011). Cell 147: 947–947.

Chen S. 2023. Ultrafast one-pass FASTQ data preprocessing, quality control, and deduplication using fastp. Imeta 2: e107.

Chen S, Zhou Y, Chen Y, Gu J. 2018. fastp: an ultra-fast all-in-one FASTQ preprocessor. Bioinformatics 34: i884–i890.

Cho JW, Cao JY, Hemberg M. 2024. Joint analysis of mutational and transcriptional landscapes in human cancer reveals key perturbations during cancer evolution. Genome Biology 25.

Clemson CM, Hutchinson JN, Sara SA, Ensminger AW, Fox AH, Chess A, Lawrence JB. 2009. An Architectural Role for a Nuclear Noncoding RNA: RNA Is Essential for the Structure of Paraspeckles. Mol Cell 33: 717–726.

Dobin A, Davis CA, Schlesinger F, Drenkow J, Zaleski C, Jha S, Batut P, Chaisson M, Gingeras TR. 2013. STAR: ultrafast universal RNA-seq aligner. Bioinformatics 29: 15–21.

Gradnigo JS, Majumdar A, Norgren RB, Jr., Moriyama EN. 2016. Advantages of an Improved Rhesus Macaque Genome for Evolutionary Analyses. PLoS One 11: e0167376.

Hao Y, Stuart T, Kowalski MH, Choudhary S, Hoffman P, Hartman A, Srivastava A, Molla G, Madad S, Fernandez-Granda C et al. 2024. Dictionary learning for integrative, multimodal and scalable single-cell analysis. Nat Biotechnol 42: 293–304.

Hebda-Bauer EK, Hagenauer MH, Munro DB, Blandino P, Jr., Meng F, Arakawa K, Stead JDH, Chitre AS, Ozel AB, Mohammadi P et al. 2025. Bioenergetic-related gene expression in the hippocampus predicts internalizing vs. externalizing behavior in an animal model of temperament. Front Mol Neurosci 18: 1469467.

Iyer MK, Niknafs YS, Malik R, Singhal U, Sahu A, Hosono Y, Barrette TR, Prensner JR, Evans JR, Zhao S et al. 2015. The landscape of long noncoding RNAs in the human transcriptome. Nat Genet 47: 199–208.

Kaur G, Perteghella T, Carbonell-Sala S, Gonzalez-Martinez J, al e. 2024. The GENCODE CLS project: massively expanding the lncRNA catalog through capture long-read RNA sequencing. bioRxiv doi:10.1101/2024.10.29.620654.

Kino T, Hurt DE, Ichijo T, Nader N, Chrousos GP. 2010. Noncoding RNA Gas5 Is a Growth Arrest- and Starvation-Associated Repressor of the Glucocorticoid Receptor. Science Signaling 3.

Kovaka S, Zimin AV, Pertea GM, Razaghi R, Salzberg SL, Pertea M. 2019. Transcriptome assembly from long-read RNA-seq alignments with StringTie2. Genome Biol 20: 278.

Lagarde J, Uszczynska-Ratajczak B, Carbonell S, Perez-Lluch S, Abad A, Davis C, Gingeras TR, Frankish A, Harrow J, Guigo R et al. 2017. High-throughput annotation of full-length long noncoding RNAs with capture long-read sequencing. Nat Genet 49: 1731–1740.

Lee JT. 2012. Epigenetic regulation by long noncoding RNAs. Science 338: 1435–1439.

Lee S, Kopp F, Chang TC, Sataluri A, Chen BB, Sivakumar S, Yu HT, Xie Y, Mendell JT. 2016. Noncoding RNA Regulates Genomic Stability by Sequestering PUMILIO Proteins. Cell 164: 69–80.

Li H, Handsaker B, Wysoker A, Fennell T, Ruan J, Homer N, Marth G, Abecasis G, Durbin R, Genome Project Data Processing S. 2009. The Sequence Alignment/Map format and SAMtools. Bioinformatics 25: 2078–2079.

Li J, Choi J, Cheng X, Ma J, Pema S, Sanes JR, Mardon G, Frankfort BJ, Tran NM, Li Y et al. 2024. Comprehensive single-cell atlas of the mouse retina. iScience 27: 109916.

Mapleson D, Venturini L, Kaithakottil G, Swarbreck D. 2018. Efficient and accurate detection of splice junctions from RNA-seq with Portcullis. Gigascience 7.

Mattick JS, Amaral PP, Carninci P, Carpenter S, Chang HY, Chen LL, Chen RS, Dean C, Dinger ME, Fitzgerald KA et al. 2023. Long non-coding RNAs: definitions, functions, challenges and recommendations. Nat Rev Mol Cell Bio 24: 430–447.

Pak M, Saurty-Seerunghen MS, Wise K, Abera TA, Lama C, Parghi N, Kang T, Sun XT, Gao Q, Bao LM et al. 2026. Co-mapping clonal and transcriptional heterogeneity in somatic evolution via GoT-Multi. Cell Genomics 6.

Pertea M, Pertea GM, Antonescu CM, Chang TC, Mendell JT, Salzberg SL. 2015. StringTie enables improved reconstruction of a transcriptome from RNA-seq reads. Nat Biotechnol 33: 290–295.

Pertea M, Shumate A, Pertea G, Varabyou A, Breitwieser FP, Chang YC, Madugundu AK, Pandey A, Salzberg SL. 2018. CHESS: a new human gene catalog curated from thousands of large-scale RNA sequencing experiments reveals extensive transcriptional noise. Genome Biol 19: 208.

Quinlan AR, Hall IM. 2010. BEDTools: a flexible suite of utilities for comparing genomic features. Bioinformatics 26: 841–842.

Rinn JL, Chang HY. 2020. Long Noncoding RNAs: Molecular Modalities to Organismal Functions. Annu Rev Biochem 89: 283–308.

Rinn JL, Kertesz M, Wang JK, Squazzo SL, Xu X, Brugmann SA, Goodnough LH, Helms JA, Farnham PJ, Segal E et al. 2007. Functional demarcation of active and silent chromatin domains in human HOX loci by noncoding RNAs. Cell 129: 1311–1323.

Ritchie ME, Phipson B, Wu D, Hu Y, Law CW, Shi W, Smyth GK. 2015. limma powers differential expression analyses for RNA-sequencing and microarray studies. Nucleic Acids Res 43: e47.

Robinson JT, Thorvaldsdottir H, Winckler W, Guttman M, Lander ES, Getz G, Mesirov JP. 2011. Integrative genomics viewer. Nat Biotechnol 29: 24–26.

Statello L, Guo CJ, Chen LL, Huarte M. 2021. Gene regulation by long non-coding RNAs and its biological functions. Nat Rev Mol Cell Bio 22: 96–118.

Thorvaldsdottir H, Robinson JT, Mesirov JP. 2013. Integrative Genomics Viewer (IGV): high-performance genomics data visualization and exploration. Brief Bioinform 14: 178–192.

Traag VA, Waltman L, van Eck NJ. 2019. From Louvain to Leiden: guaranteeing well-connected communities. Sci Rep 9: 5233.

Trapnell C, Roberts A, Goff L, Pertea G, Kim D, Kelley DR, Pimentel H, Salzberg SL, Rinn JL, Pachter L. 2012. Differential gene and transcript expression analysis of RNA-seq experiments with TopHat and Cufflinks. Nat Protoc 7: 562–578.

Tripathi V, Ellis JD, Shen Z, Song DY, Pan Q, Watt AT, Freier SM, Bennett CF, Sharma A, Bubulya PA et al. 2010. The Nuclear-Retained Noncoding RNA MALAT1 Regulates Alternative Splicing by Modulating SR Splicing Factor Phosphorylation. Mol Cell 39: 925–938.

Weisman CM. 2022. The Origins and Functions of De Novo Genes: Against All Odds? J Mol Evol 90: 244–257.

Weisman CM, Murray AW, Eddy SR. 2020. Many, but not all, lineage-specific genes can be explained by homology detection failure. PLoS Biol 18: e3000862.

Weisman CM, Murray AW, Eddy SR. 2022. Mixing genome annotation methods in a comparative analysis inflates the apparent number of lineage-specific genes. Curr Biol 32: 2632–2639 e2632.

